# PU.1-driven enrichment enables microglia profiling from frozen brain tissue using the high-throughput Smart-seq3xpress method

**DOI:** 10.1101/2025.08.11.669607

**Authors:** Dominika Dostalova, Pavel Abaffy, Eva Rohlova, Jan Kriska, Tomas Knotek, Jana Tureckova, Denisa Kirdajova, Miroslava Anderova, Lukas Valihrach

## Abstract

Single-cell transcriptomics has revealed the central role of microglia in brain development, homeostasis, and disease, particularly in the context of neuroinflammation. While single-cell RNA-sequencing enables targeted microglial analysis from fresh tissue, studying these cells in cryopreserved or archival samples remains challenging due to the lack of protocols for their specific enrichment.

We introduce a method for the selective isolation of microglial nuclei from fresh-frozen brain tissue using the transcription factor PU.1 as a nuclear marker. To stabilize PU.1 for reliable detection, a brief formaldehyde fixation step is applied. The protocol is fully compatible with Smart-seq3xpress, a high-sensitivity, full-length transcriptomic method offering isoform- and allele-level resolution, making the workflow scalable and cost-efficient. We benchmarked the method in a mouse model of ischemic stroke, evaluating both technical performance and its ability to capture biologically meaningful microglial states. Compared to standard single-nucleus protocols, our approach yielded higher gene and UMI counts and a greater proportion of coding reads. Transcriptomic profiles closely matched those from whole-cell RNA-sequencing including the detection of activation markers and diverse microglial subpopulations.

This approach addresses key limitations of single-nucleus RNA - sequencing and opens new possibilities for studying microglial states in cryopreserved and archival brain tissue, broadening access to cellular insights in both basic and translational research.

## INTRODUCTION

Microglia, the resident immune cells of the central nervous system, play essential roles in brain development, homeostasis, and pathology. Single-cell (scRNA-seq) and single-nucleus RNA sequencing (snRNA-seq) have significantly advanced the capacity to study microglial heterogeneity and function (1). However, scRNA-seq requires fresh tissue, limiting its use for postmortem or biobanked human samples, which are predominantly available as frozen or fixed material.

snRNA-seq provides access to such preserved tissues but remains suboptimal for studying microglia. A common enrichment strategy in scRNA-seq relies on the surface marker CD11b, but this cannot be used for nuclei-based workflows, as nuclear membranes lack surface proteins. Without nuclear-specific markers, enrichment is not possible, resulting in data dominated by other cell types. As a result, snRNA-seq datasets often contain too few microglia for robust analysis. This increases sequencing costs and reduces resolution, especially for rare or transient microglial states. Additionally, recent work has shown that snRNA-seq underrepresents many transcripts associated with microglial activation, including key immune and disease-related genes, calling into question its sensitivity for capturing cellular responses in pathology (2).

To overcome these limitations, we developed a protocol that enables selective enrichment of microglial nuclei using PU.1, a transcription factor localized to the nucleus and specific to the myeloid lineage (3). A brief formaldehyde fixation step stabilizes PU.1 detection (4), and the protocol is made compatible with full-length RNA sequencing (Smart-seq3xpress) through the incorporation of thermolabile Proteinase K for effective decrosslinking while preserving RNA integrity (5,6). Smart-seq3xpress combines high sensitivity with full-length transcript coverage, enabling isoform-level resolution and allele-specific analyses at single-cell depth (7).

We benchmarked this protocol in a mouse model of ischemic stroke (8), a condition characterized by pronounced microglial activation, and show that our approach robustly captures both homeostatic and activated microglial states. By increasing microglial representation and preserving activation-related signatures, the protocol enhances both the technical performance and biological interpretability of snRNA-seq datasets. Its compatibility with archived tissue makes it broadly applicable to studies of neuroinflammation, neurodegeneration, and other brain disorders where microglia play a critical role.

## RESULTS

### PU.1-based enrichment enhances snRNA-seq profiling of microglia in frozen brain tissue

We developed a protocol for the selective enrichment of microglial nuclei from fresh-frozen brain tissue using the transcription factor PU.1 as a nuclear marker. To preserve the PU.1 epitope for reliable detection, a brief formaldehyde fixation step was introduced, followed by nuclei isolation under mild conditions (FixedNuclei). The protocol was optimized for compatibility with full-length single-cell transcriptomic profiling using Smart-seq3xpress (7), including an additional lysis step with thermolabile proteinase K (6) to reverse crosslinking while preserving RNA integrity **(Methods).**

To evaluate this approach, we applied it to cortical tissue from a mouse model of focal cerebral ischemia middle cerebral artery occlusion (MCAO), collected 7 days post-stroke, along with sham-operated controls. We compared the transcriptomic profiles generated using the FixedNuclei protocol to two established methods: (1) whole-cell single-cell RNA sequencing (LiveCells), in which dissociated cells are enriched using CD11b-based magnetic selection to isolate microglia and related myeloid cells; and (2) single-nucleus RNA sequencing of unfixed nuclei (LiveNuclei), in which nuclei were isolated from the CD11b⁺-enriched cell suspension. While CD11b is widely used for isolating myeloid cells in live-cell workflows, it cannot be used for direct nuclear enrichment. In contrast, the FixedNuclei protocol enables targeted sorting of PU.1⁺ nuclei from lightly fixed, frozen tissue, allowing for selective microglial enrichment in a nuclei-compatible format (**Fig. 1A**).

**Fig. 1:**
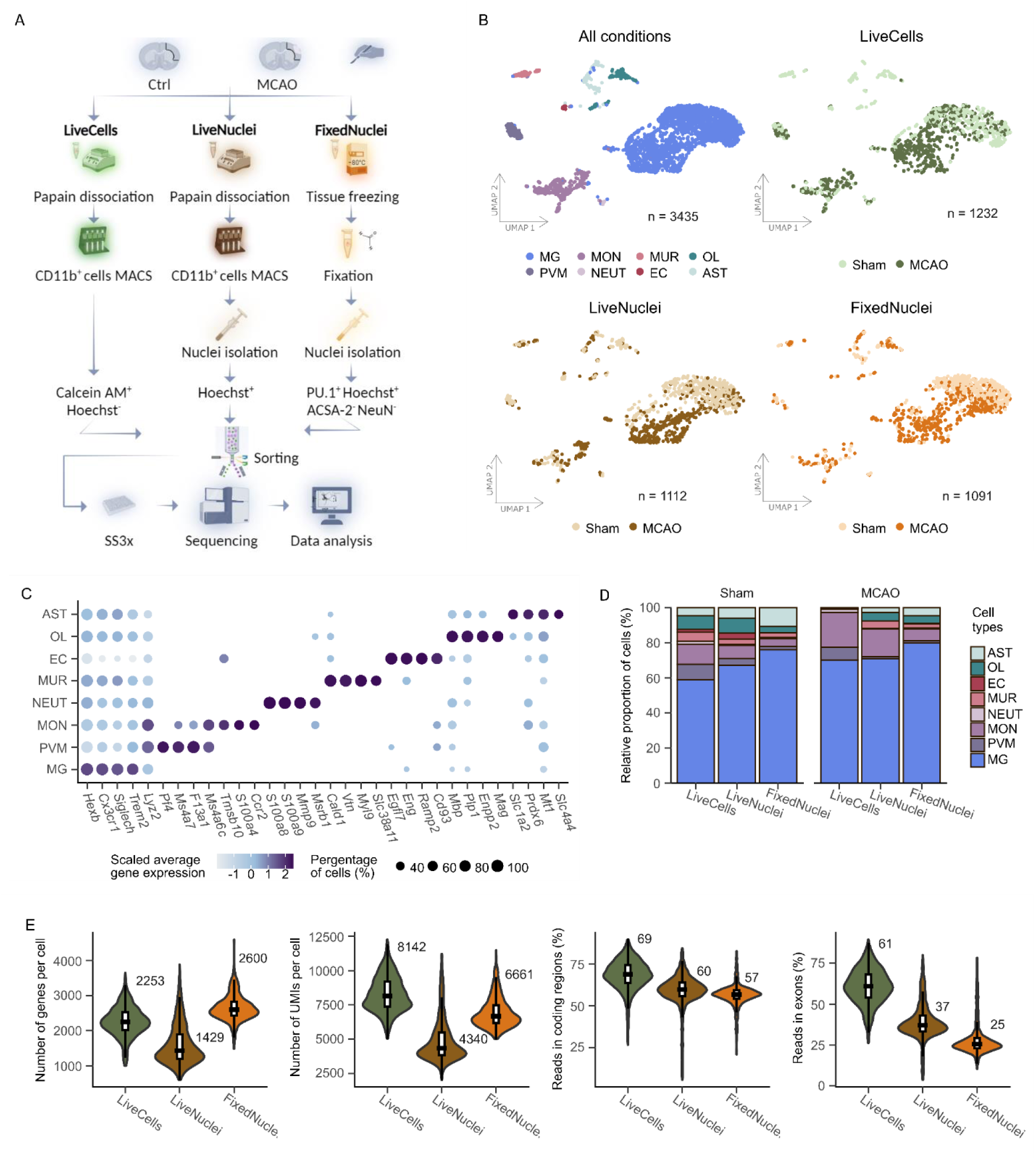
Comparison of three enrichment protocols for microglia populations. **A** Overview of the experimental pipeline: Tissue was processed using three protocols (LiveCells, LiveNuclei, FixedNuclei) for the enrichment of microglia populations. Isolated cells were sorted into 384-well plates for RNA-seq library preparation using the Smart-seq3xpress approach, followed by downstream analysis (Created with BioRender.com). **B** UMAP visualization showing identified clusters across all conditions: microglia (MG, n = 2 420), monocytes (MON, n = 387), perivascular macrophages (PVM, n = 144), oligodendrocytes (OL, n = 162), astrocytes (AST, n = 156), mural cells (MUR, n = 97), epithelial cells (EC, n = 36), and neutrophils (NEUT, n = 33). Cluster representation across isolation protocols (LiveCells, LiveNuclei, and FixedNuclei) and conditions (Sham vs MCAO) is included. **C** Dot plot showing canonical marker genes used for cell cluster identification. **D** Stacked bar plot representing the relative proportions of cells across isolation protocols (LiveCells, LiveNuclei, and FixedNuclei) divided into MCAO and Sham groups. **E** Violin plots with overlaid box plots summarizing technical parameters: number of genes per cell, number of UMIs per cell, reads in coding regions and reads in exons for LiveCells, LiveNuclei, and FixedNuclei. Median values are displayed in the top-right corner of each plot. Box plots show medians (black line), first and third quartiles (white boxes), and whiskers extending to 1.5× the interquartile range.

Across all protocols, we identified eight major brain cell types: microglia (MG), monocytes (MON), perivascular macrophages (PVM), oligodendrocytes (OL), astrocytes (AST), mural cells (MUR), neutrophils (NEUT), and epithelial cells (EC) (**Fig. 1B**). Microglia were characterized by high expression of canonical markers including *Hexb*, *Cx3cr1*, *Siglech*, and *Trem2* (**Fig. 1C**), and were the dominant population, comprising over 60% of the dataset (2,420 cells/nuclei) (**Fig. 1D**).

To enrich microglia, we employed magnetic sorting (MACS) based solely on CD11b expression, a commonly used myeloid marker. Unlike the CD11b⁺/CD45^low^ gating strategy often applied in FACS, our simplified approach reduces cell stress and enhances viability, helping preserve microglial transcriptomic integrity (9,10). However, CD11b is also expressed by other myeloid cells, which likely contributed to the presence of monocytes and perivascular macrophages in the LiveCells and LiveNuclei protocols. In contrast, the FixedNuclei protocol used a more selective nuclear marker-based strategy for sorting, resulting in higher microglial purity and reduced contamination from other lineages (**Fig. 1D**).

We next evaluated protocol performance based on key technical parameters. Surprisingly, the FixedNuclei protocol detected the highest number of genes per cell (2,600), surpassing both LiveCells (2,253) and LiveNuclei (1,429). Similarly, it yielded more UMIs per cell (6,661) than LiveNuclei (4,340), though not as many as LiveCells (8,142) (**Fig. 1E**). These trends were consistent across both MCAO and Sham conditions **(SFig. 1A)** and were largely reproduced in an independent experiment, except for slightly lower gene counts per cell in the FixedNuclei protocol **(SFig. 1B).** All comparisons were performed at a standardized sequencing depth of 20,000 reads per cell to maintain analytical consistency. At higher read depths (5×10⁴, 7.5×10⁴, and 1×10⁵), performance gaps became more pronounced, with no plateau observed (**SFig. 1C**), indicating true differences in RNA capture efficiency rather than coverage bias.

Read mapping statistics further reflected expected differences between nuclear and whole-cell preparations. For coding regions, FixedNuclei achieved 57% of reads mapped—slightly lower than LiveNuclei (60%) and below LiveCells (69%). Reads mapping to exonic regions followed a similar trend: 25% for FixedNuclei, 37% for LiveNuclei, and 61% for LiveCells **(Fig. 1E)**. These differences are expected, as nuclei contain higher levels of pre-mRNA and retained introns, whereas cytoplasmic RNA is enriched for fully spliced, exon-dense transcripts (11).

Together, these results demonstrate that our PU.1-based enrichment strategy preserves RNA integrity and improves the quality of nuclei-based transcriptomic data compared to standard snRNA-seq. While whole-cell protocols still offer the richest transcriptomic coverage, the FixedNuclei protocol enables robust profiling of microglial populations from frozen tissue— addressing the key limitations of marker availability, data sparsity, and RNA degradation in archival samples.

### PU.1-based approach preserves microglial heterogeneity in frozen tissue

To assess how well microglial diversity is preserved across protocols, we performed unsupervised clustering of all microglia and analysed the resulting transcriptional states. This allowed us to determine whether our PU.1-based FixedNuclei protocol captures biologically meaningful microglial heterogeneity comparable to established methods.

We identified six microglial subpopulations across all datasets: two homeostatic populations (HM1, HM2), activated response microglia (ARM), interferon response microglia (IRM), an intermediate state (INTER), and a small population of dividing microglia (DIV) (**Fig. 2A**). HM1 and HM2 expressed canonical homeostatic genes such as *P2ry12* and *Tmem119*, with HM2 also showing elevated levels of *Hexb*, *C1qa*, *C1qb*, and *Trem2*. These populations were most abundant in sham samples but also remained detectable in post-stroke conditions. In contrast, ARM, IRM, and INTER subtypes were predominantly observed in MCAO samples. ARM cells expressed activation-associated genes including *Lyz2*, *Apoe*, *Cst7*, *Lpl*, *Clec7a*, *Ccl6*, *Ccl3*, and *Ccl4*. IRM cells were characterized by interferon response markers such as *Ifit3*, *Irf7*, and *Ifitm3*. The INTER population exhibited reduced homeostatic marker expression but lacked strong upregulation of activation markers (**Fig. 2B–C**). We also note occasional detection of microglial marker genes in non-microglial clusters, which is likely a result of index hopping—a known technical artifact in single-cell sequencing that can lead to barcode misassignment, particularly when one cell type dominates the dataset (12).

**Fig. 2:**
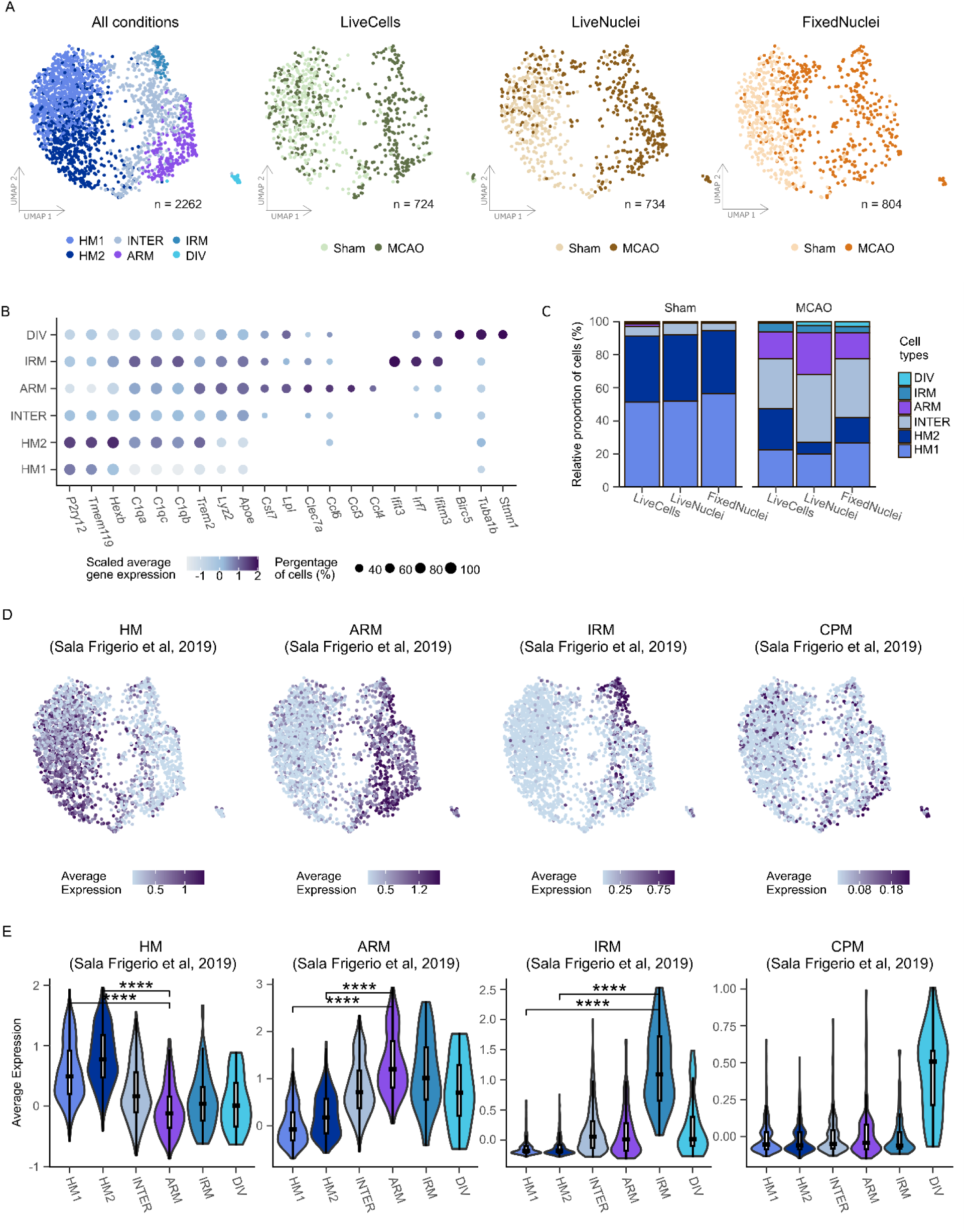
Characterization of microglia subpopulations. **A** UMAP visualization showing identified microglia clusters across all conditions, including homeostatic 1 (HM1, n = 841), homeostatic 2 (HM2, n = 609), intermediate (INTER, n = 489), activated response (ARM, n = 238), interferon response (IRM, n = 56), and dividing (DIV, n = 29) microglia. Additionally, the cluster representation is displayed for each isolation protocol (LiveCells, LiveNuclei, and FixedNuclei) in both Sham and middle cerebral artery occlusion (MCAO) conditions. **B** Dot plot showing canonical marker genes used for the identification of microglia subclusters. **C** Stacked bar plot representing the relative proportions of cells across isolation protocols (LiveCells, LiveNuclei, and FixedNuclei) divided into MCAO and Sham groups. **D** Feature plot visualizing the average expression of microglia marker genes as defined by Sala Frigerio et al. (13) - homeostatic (HM), ARM, IRM and cycling/proliferating (CPM) microglia. **E** Violin plots with overlaid box plots summarizing the average expression of microglia marker genes as defined by Sala Frigerio et al. (13) across our microglia subpopulations. Box plots show medians (black line), first and third quartiles (white boxes), and whiskers extending to 1.5× the interquartile range. Statistical significance of differences in average gene expression between HM1, HM2 vs ARM populations was evaluated using an unpaired Wilcoxon rank-sum test.

To validate these classifications, we compared our cluster-specific gene signatures with those reported by Sala Frigerio et al. in a mouse model of Alzheimer’s disease (13). In their study, ARM and IRM states emerged gradually over several months in response to amyloid-β accumulation. In contrast, we observed similar microglial states within just seven days after stroke, highlighting the rapid response of microglia to acute injury. Marker genes from that study were highly enriched in our corresponding clusters, supporting the robustness of our classification (**Fig. 2D–E**).

In summary, we identified six transcriptionally distinct microglial subpopulations across all protocols, capturing both homeostatic and activated states. All subtypes were consistently detected across protocols, demonstrating the preservation of microglial identity and transcriptional heterogeneity in our dataset. This highlights the robustness of our classification and confirms that the PU.1-based FixedNuclei method enables reliable profiling of microglia from frozen tissue.

### PU.1-based protocol enables detection of inflammatory microglial states

As key mediators of neuroinflammation, ARM are frequently studied in the context of neurological diseases. However, there is ongoing discussion regarding how isolation methods affect the detection of activation-related transcriptional changes (2). To address this, we compared the performance of our PU.1-based FixedNuclei protocol with the widely used LiveCells and LiveNuclei approaches.

We first identified differentially expressed genes (DEGs) between homeostatic microglia (combination of HM1 and HM2; HM) and ARM populations across all conditions (**Fig. 3A**). A shared set of 89 DEGs was found in the ARM population across all three protocols (**Fig. 3B**), indicating that a common activation signature is consistently captured, regardless of the isolation method. This gene set includes canonical activation markers such as *Apoe*, *Lyz2*, *Cd63*, and *Clec7a*, associated with phagocytosis and immune activation, as well as *Lpl*, *Ctsb*, *Hif1a*, and *Ccl3*, which reflect metabolic adaptation and inflammatory signalling in activated microglia.

**Fig. 3:**
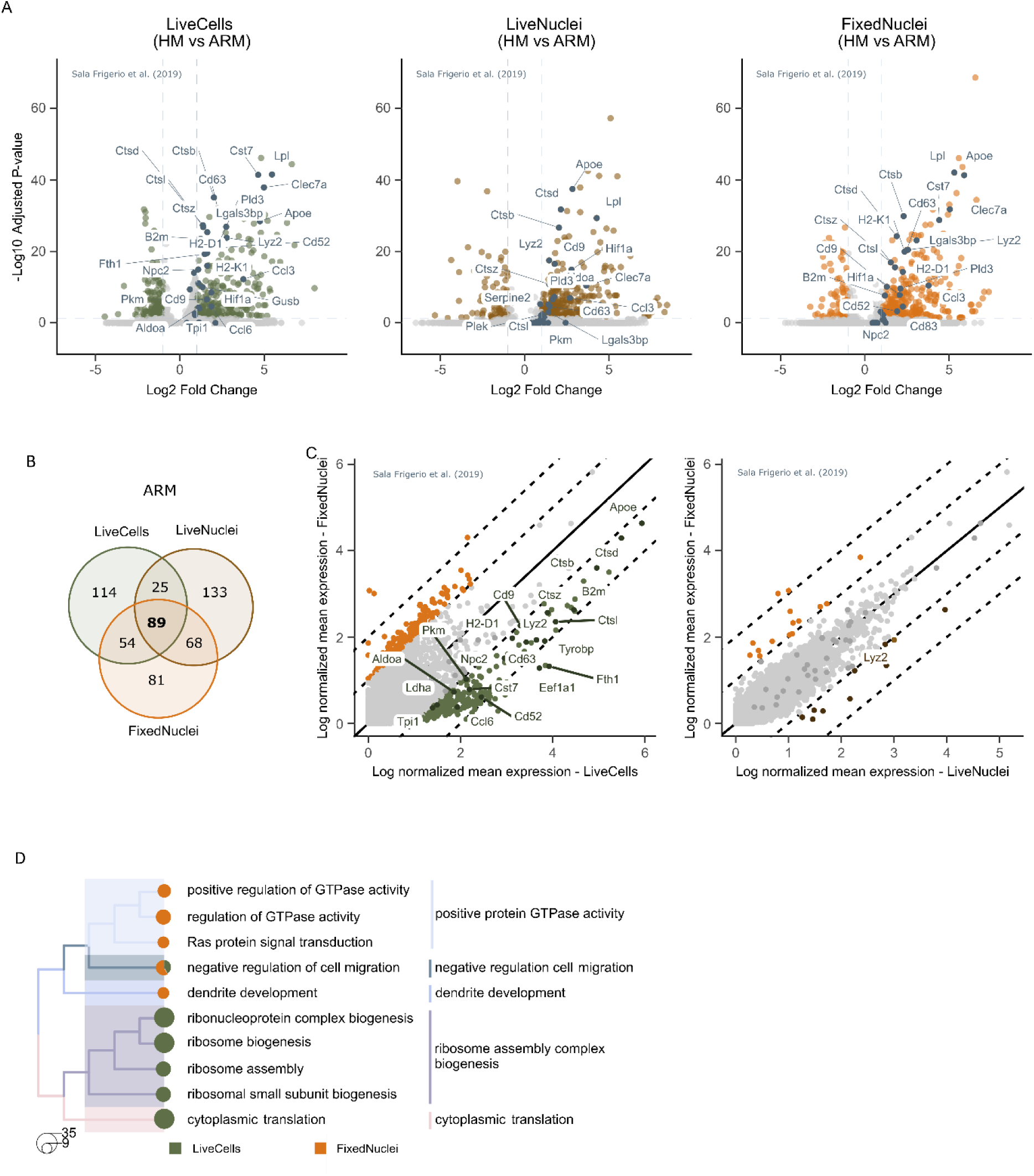
Comparison of microglia activation over isolation protocols. **A** Volcano plots illustrating differential gene expression between homeostatic microglia (combination of HM1 and HM2; HM) and ARM over isolation protocols. Each point represents the average value of a transcript. Colour (green, brown or oranges) points indicate transcripts with a log2 fold change > 1 and an adjusted p-value > 0.05 (−log10 scale). Highlighted dots (dark grey) represent markers of the ARM population, as characterized by Sala Frigerio et al. (13). **B** The Venn diagram illustrates the overlap of DEGs in the ARM population across the different isolation protocols. DEGs were selected based on a log2 fold change > 1 and an adjusted p-value < 0.05. **C** Scatter plots showing the mean normalized gene abundance in the ARM population, comparing isolation protocols: LiveCells vs. FixedNuclei and LiveNuclei vs. FixedNuclei. The solid black line represents no fold change, while the dotted black lines denote 2- and 4-fold differences between isolation protocols. Genes labeled in the area with >1 log-normalized mean expression are associated with microglial activation, as characterized by Sala Frigerio et al. (13). Full results are provided (**STab. 4–5**). **D** Tree plot of the top 10 significant GOBP terms identified by overrepresentation analysis, grouped into five clusters. The analysis includes genes with >1 log-normalized mean expression identified from the differential expression analysis LiveCells vs FixedNuclei (Fig. 3C). Full results are provided (**STab. 6**).

To assess how activation gene expression levels varied across protocols, we examined log₂ fold changes in ARM and visualized mean gene expression using scatter plots (**Fig. 3C**). ARM-associated genes showed generally higher expression in LiveCells compared to FixedNuclei, particularly for inflammatory and phagocytic markers. In contrast, differences between FixedNuclei and LiveNuclei were smaller, suggesting closer alignment between the nuclear protocols. To explore the functional relevance of these differences, we performed Gene Ontology (GO) analysis on genes enriched in each protocol. LiveCells showed upregulation of ribosome-related pathways, while FixedNuclei was enriched for processes such as GTPase regulation and dendrite development (**Fig. 3E**). This pattern aligns with findings by Thrupp et al., who reported reduced detection of activation genes and translational machinery in nuclei compared to whole cells (2).

In summary, our PU.1-based FixedNuclei protocol effectively detects activated microglial states in frozen brain tissue. Although expression levels differ from whole-cell data due to the absence of cytoplasmic RNA, activated microglial gene programs—including those relevant to inflammation and disease—can be reliably studied using this approach.

### PU.1-based profiling captures cell–cell interactions among microglia

While the transcriptional activation of microglia in brain pathologies is well-documented, analysing intercellular communication provides additional insights into how cell populations interact and coordinate their responses during the disease progression. In this study, we focused on communication specifically among microglial populations. Due to the low number of cells captured in the LiveCells protocol, the DIV cluster was excluded from analysis.

We assessed communication using two quantitative metrics: the number of inferred ligand–receptor interactions and overall interaction strength. Both LiveCells and FixedNuclei protocols yielded a similar number of interactions (155 vs. 163), whereas LiveNuclei detected only 48 interactions (**Fig. 4A, left**). A comparable pattern was observed for interaction strength: LiveCells and FixedNuclei reached similar values (0.239 and 0.232), while LiveNuclei showed a fivefold lower value (0.04) (**Fig. 4A, right**). These trends were further illustrated in chord diagrams, which showed denser and more evenly distributed interactions in LiveCells and FixedNuclei compared to LiveNuclei (**Fig. 4B**).

**Fig. 4:**
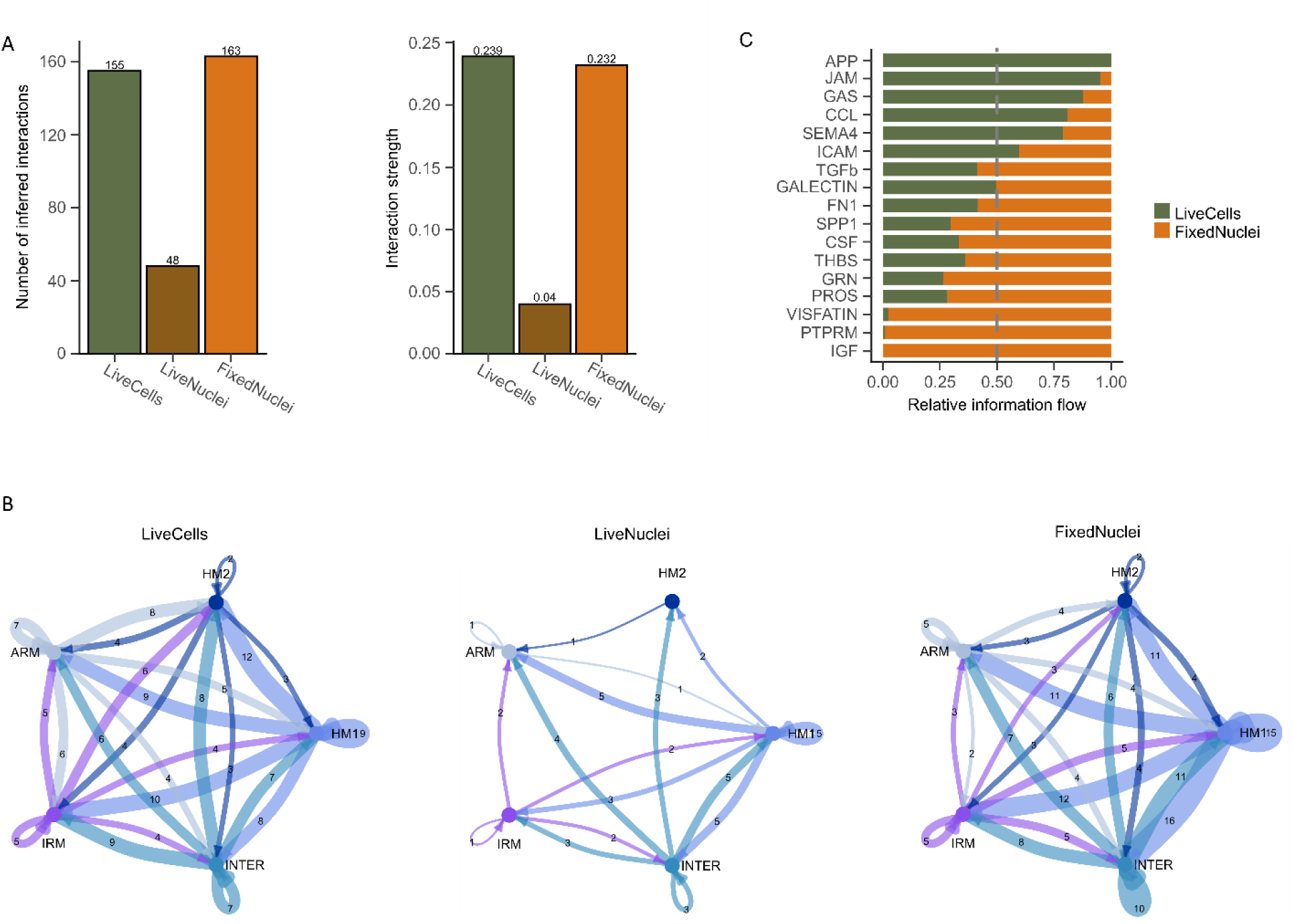
Cell–cell communication in microglial subclusters across isolation protocols. **A** (**left**) Number of predicted cell–cell interactions across different isolation protocols. **A** (**right**) Strength of predicted cell–cell interactions across different isolation protocols. **B** The number of significant ligand-receptor pairs between each pair of cell populations is represented, with edge width proportional to the total number of detected interactions. Thicker edges indicate a higher number of ligand-receptor pairs between the connected populations. **C** Significant signalling pathways were ranked based on differences in overall information flow, calculated by summing communication probabilities, to compare cell–cell communication across isolation protocols.

Given the poor performance of the LiveNuclei dataset, we excluded it from downstream pathway-level comparisons. Among the remaining protocols, we identified 17 signalling pathways, 15 of which were shared between LiveCells and FixedNuclei. Analysis across all identified pathways revealed expected components of microglial response to stroke, such as inflammatory pathways (e.g. CCL, ICAM, GAS) (14,15), clearance signals (e.g. SPP1, GRN) (16,17), and repair-associated signalling pathways that promote anti-inflammatory phenotypes (e.g., TGF-β, THBS) (18,19), tissue remodelling (e.g., FN1) (20), and functional recovery (e.g. GRN, IGF) (21,22). However, the relative information flow of these pathways varied between protocols (**Fig. 4C**), with several—including IGF, PTPRM, and VISFATIN—showing predominant detection in FixedNuclei, while pathways such as APP, JAM, and GAS were more prominent in LiveCells. This variation may reflect both technical biases and biological variability in the MCAO model.

In conclusion, the PU.1-based FixedNuclei protocol effectively captures microglial cell–cell communication networks and signalling dynamics, producing results that closely resemble those obtained from whole-cell protocols. Despite the inherent limitations of snRNA-seq—such as reduced detection of certain ligands or receptors—the PU.1-based approach enables meaningful inference of intercellular signalling and is well suited for studies using frozen or archived brain tissue.

## DISCUSSION

Understanding microglial heterogeneity in pathological contexts requires not only sensitive transcriptomic techniques but also reliable enrichment strategies that preserve cellular diversity. Although CD11b is commonly used to enrich microglia from fresh tissue, its application to frozen samples has also been reported (23). However, this approach often results in poor RNA quality, likely due to the challenges of processing fixed cell suspensions from thawed postmortem tissue. To address these limitations, we introduce a PU.1-based protocol optimized for the isolation and transcriptomic profiling of microglia from fresh frozen brain tissue, providing a more reliable approach for studying these cells under disease conditions.

We selected PU.1 as the primary nuclear marker for microglial enrichment due to its stable expression across both homeostatic and reactive states. As a master regulator of microglial identity and function, PU.1 represents a more stable and reliable marker compared to alternatives like SALL1 and IRF8, whose expression fluctuates with activation status. Specifically, SALL1 is often downregulated in activated microglia (24), whereas IRF8 is typically upregulated (25).

Our approach is supported by previous findings from Alzheimer’s disease studies, where PU.1-based enrichment was successfully applied without fixation, allowing effective sorting and transcriptomic analysis (26,27). The use of protease inhibitors during nuclei isolation in those protocols likely contributed to improved epitope preservation and antibody binding. Future comparisons between fixation and protease inhibition could help optimize epitope stability and overall protocol robustness.

Our protocol employed protein-based enrichment by targeting the nuclear transcription factor PU.1; alternatively, microglial enrichment can be achieved using RNA-based approaches. One such method is nuclampFISH, an amplified RNA FISH approach that enables cell sorting based on nuclear RNA expression and has been applied in downstream chromatin profiling (28). Another example is, HCR-FlowFISH, which uses hybridization chain reaction amplification to detect specific transcripts and has been successfully applied to nuclei from frozen tissue in the context of PERFF-seq for enriching rare cell populations prior to single-cell RNA sequencing (29). RNA-based enrichment is more labour-intensive and time-consuming than our approach because it requires hybridization and signal-amplification steps. However, it offers boarder flexibility, as probe design can be tailored to selectively enrich the specific cell population of interest.

Moreover, the use of Smart-seq3xpress provided high-sensitivity and full-length transcript coverage, offering a significant advantage over 10x Genomics-based approaches. While we did not fully leverage the isoform-level resolution in this analysis—to ensure comparability with previous studies—Smart-seq3xpress remains a powerful platform for future investigations requiring greater transcriptomic depth, such as alternative splicing or low-abundance gene detection.

Despite these advances, our study has limitations. Biological variability introduced by the MCAO model may influence gene expression patterns, and the relatively low number of microglial cells and nuclei analysed (2,262 total across three protocols) may limit the statistical power and depth of downstream analyses, increasing the potential for bias in interpretation.

In conclusion, our PU.1-based protocol offers a reliable method for enriching microglia from frozen brain samples, enabling high-quality transcriptomic profiling even in pathological conditions. This approach lays the groundwork for future studies aiming to resolve microglial diversity and function at single-cell resolution using full-length transcriptomic methods.

## CONCLUSION

In summary, we present a PU.1-based protocol for enriching microglia from fresh-frozen brain tissue, optimized for high-sensitivity transcriptomic analysis using Smart-seq3xpress. By targeting a nuclear marker with stable expression across microglial states, we achieved consistent enrichment suitable for profiling both homeostatic and activated populations. While our analysis was limited by biological variability inherent to the MCAO model and a modest number of captured cells, the results demonstrate the feasibility and reliability of this approach. This protocol provides a robust foundation for future studies of microglial heterogeneity in pathological conditions and offers a path toward leveraging full-length transcriptomic data to get deeper insights into microglial function and gene regulation.

## DECLARATIONS

### Ethics approval and consent to participate

All procedures involving the use of laboratory mice were performed in accordance with the European Communities Council Directive (November 24, 1986; 86/609/EEC) and the Animal Welfare Guidelines approved by the Institute of Experimental Medicine of the Czech Academy of Sciences (approval numbers 50/2020 and 112-2023-P).

### Consent for publication

Not applicable.

### Availability of data and materials

The data that support the findings of this study are publicly available Zenodo and the code for analysis is available on [GitHub](https://github.com/GliaOmicsLab/microglia-sc-snRNAseq). Data from the main experiment ([all cells](https://scarfweb.nygen.io/eu-central-1/public/q2acwcw8) and the [subset of microglia](https://scarfweb.nygen.io/eu-central-1/public/wxvqmskh)) are available for online analysis via the Nygen portal. A detailed version of the optimized protocol is available at Protocols.io.

### Authors’ contributions

DD and PA designed the research, isolated and processed samples, and conducted data analysis. DD, ER, and PA optimized the Smart-seq3xpress protocol. DD wrote the manuscript, while TK and DK performed surgical procedures. JK and JT were responsible for mouse breeding, managed mouse model work, and isolated mouse brains. MA and LV coordinated the overall project and reviewed the manuscript.

## Acknowledgements

We would like to thank Martin Pavlíček for his technical assistance in the laboratory. We acknowledge the GeneCore Facility in BIOCEV for their help with library preparation and the Imaging Methods Core Facility, supported by the Ministry of Education, Youth and Sports of the Czech Republic (LM2023050 Czech-Bioimaging), for their help with cell sorting. We would also like to acknowledge the use of Biorender.com for creating the schematic illustration included in this article and ChatGPT for improving readability of text.

## METHODS

### Experimental model - middle cerebral artery occlusion (MCAO) model

Experiments were performed on 22 to 29-week-old C57Bl/6J male mice (Jackson Laboratory, #000664). The mice were housed under a 12-hr light/dark cycle with ad libitum access to food and water. The method for inducing focal cerebral ischemia by permanent middle cerebral artery occlusion (MCAO) has been detailed previously (8). Briefly, the middle cerebral artery was occluded at the distal segment using bipolar tweezers. Sham surgery was used as a control, and our previous results showed that sham surgery did not impact gene expression compared to naïve controls (8).

### Isolation of cortical tissue from the mouse brain

Deeply anesthetized mice with sodium pentobarbital (PTB; 100 mg/kg, i.p.; Sigma-Aldrich) were perfused transcardially with cold (4–8°C) isolation buffer (IB) composed of (in mM): NaCl 136.0, KCl 5.4, HEPES 10.0, glucose 5.5, with a pH of 7.4 and an osmolality of 290 ± 3 mOsmol/kg. Cortical tissue was isolated from the left hemisphere. The brain (+2 mm to -2 mm from bregma) was cut, and the sham or post-ischemic parietal cortex was carefully dissected, avoiding the ventral white matter tracts. Two animals were used for each experimental condition.

### Dissociation of brain tissue into

#### a) Single-cell suspension (LiveCells protocol)

The freshly dissected cortex was placed in 1 mL of papain solution (20 U/mL) containing 60 µL of Deoxyribonuclease I (DNase I; 240 U) (both from Worthington Biochemical Corp.) and Actinomycin D (ActD; 25 µg/mL, dissolved in DMSO, both from Sigma-Aldrich). The tissue suspension was incubated at 37°C for 30 minutes with shaking at 1,000 rpm. During incubation, the tissue was triturated every 10 minutes by pipetting, first with a wide-bore P1000 tip (Axygen), followed by a standard P1000 filter tip (Axygen). After incubation, the suspension was centrifuged at 500 × g for 5 minutes at 2°C, with the brake turned off and minimal acceleration.

The supernatant was carefully removed, and the pellet was resuspended in 1 mL of ice-cold resuspension buffer consisting of IB supplemented with ovomucoid inhibitor (1 mg/mL), albumin (1 mg/mL), DNase I (200 U), and ActD (2.5 µg/mL). The suspension was incubated for 5 minutes on ice. Subsequently, 1 mL of the resuspended cells was gently layered over 3 mL of an ovomucoid protease inhibitor solution prepared in sterile Earle’s Balanced Salt Solution (EBSS) containing ovomucoid inhibitor (10 mg/mL) and albumin (10 mg/mL) (all three from Worthington Biochemical Corp). The samples were centrifuged at 200 × g for 7 minutes at 2°C.

The resulting cell pellet was resuspended in 1 mL of preservation buffer (PB), consisting of 1% bovine serum albumin (BSA; ThermoFisher Scientific) in IB supplemented with ActD (2.5 µg/mL). The resuspended cells were then filtered through a 50 µm CellTrics filter (Sysmex) and centrifuged at 300 × g for 5 minutes at 2°C.

The supernatant was removed, leaving approximately 90 µL of the cell pellet in PB. Next, 10 µL of magnetically labeled anti-CD11b antibody (1:10 dilution, Miltenyi Biotec) and 0.5 µL of Calcein Green (1:1000 dilution, ThermoFisher Scientific) were added to the suspension, and the mixture was incubated at 4°C for 15 minutes. After incubation, 1 mL of PB was added, and the sample was centrifuged at 300 × g for 5 minutes at 2°C.

The supernatant was carefully removed, and the cell pellet was resuspended in 0.5 mL of cold PB before being transferred onto a 50 µm CellTrics filter placed on an MS magnetic column positioned on OctoMACS™ Separator (both from Miltenyi Biotec). The column was washed four times with 500 µL of PB. After washing, the column was removed from the magnet, and cells were eluted using 2 × 1 mL of PB containing RNAseOUT (0.2 U/µL; ThermoFisher Scientific).

Shortly before sorting, 1 µL of Hoechst 33258 was added to the suspension. Calcein Green^+^/Hoechst 33258^-^ cells were sorted into 384-well plates containing lysis buffer (see Generation of Smart-seq3xpress libraries) (**SFig. 2A**). All sorting was performed using a BD FACSMelody system (BD FACSChorus v1.3 software) equipped with a 100-μm nozzle, with sample and plate cooling maintained at 4°C and index sorting enabled. Immediately after sorting, the plates were promptly centrifuged and stored at −80°C.

#### b) Single-nucleus suspension (LiveNuclei protocol)

The CD11b⁺ enriched single-cell population was centrifuged at 300 × g for 5 minutes at 2°C and resuspended in 0.75 mL of lysis mix (10 mM NaCl, 3 mM MgCl₂, 0.10% [v/v] NP-40, 0.32 M sucrose, 1× cOmplete Mini protease inhibitor [Roche], 10 mM Tris-HCl, pH 7.4, 0.2 U/mL RNaseOUT). The suspension was transferred to a pre-chilled Dounce tissue grinder (Sigma-Aldrich), and nuclei were released using five strokes with Pestle B.

The suspension was filtered through a 30 µm CellTrics filter into a 2 mL LoBind tube (Eppendorf). The tissue grinder was rinsed with an additional 0.75 mL of lysis mix, and the wash was filtered into the same tube. The resulting suspension was carefully layered onto 0.5 mL of 1.2 M sucrose solution (containing 0.2 U/mL RNAseOUT; prepared in 10 mM NaCl, 3 mM MgCl₂, 10 mM Tris-HCl, pH 7.4) and centrifuged at 5,000 × g for 20 minutes at 2°C. The supernatant (both phases) was removed, and the pellet was resuspended in 1 mL nuclei wash buffer (PBS-/- supplemented with 0.5% BSA and 0.2 U/mL RNAseOUT). The suspension was filtered through a 30 µm CellTrics filter.

Before sorting, 1 µL of Hoechst 33258 was added to the suspension, and Hoechst 33258^+^ nuclei were sorted into 384-well plates containing lysis buffer (see Generation of Smart-seq3xpress libraries) using fluorescence-activated nuclei sorting (FANS) under the same parameters as described above (see Single-cell suspension, LiveCells protocol) (**SFig. 2B**).

#### c) Fixed single-nucleus suspension (FixedNuclei protocol)

Freshly frozen brain cortex tissue was placed into 0.3 mL of chilled fixation solution (1% formaldehyde in PBS−/−) in a 1.5 mL LoBind tube (Eppendorf) and disrupted using a microtube pestle. The homogenized tissue was then transferred using a P1000 wide-bore tip into a 2 mL tube (Eppendorf) containing 1.6 mL of fixation solution and shaken on ice at 300 rpm for 10 minutes.

Next, 0.1 mL of 2.5 M glycine was added to the suspension, which was shaken again for an additional 5 minutes, followed by centrifugation at 1,100 × g for 5 minutes at 2°C. The supernatant was carefully removed, and the tissue pellet was resuspended in 1 mL of ice-cold lysis solution (lysis mix without 1× cOmplete Mini protease inhibitor; see Single-nucleus suspension, LiveNuclei protocol) using P1000 wide-bore pipette tips and centrifuged again at 1,100 × g for 5 minutes at 2°C. This wash step was repeated a second time to ensure thorough removal of residual fixation solution.

After the final centrifugation, the supernatant was removed, and the tissue pellet was resuspended in 1 mL of ice-cold lysis solution using P1000 wide-bore tips and transferred into a pre-chilled Dounce tissue grinder. Nuclei isolation was performed on ice with 5 strokes using Pestle A, followed by 10 strokes with Pestle B. The resulting suspension was filtered through a 30 µm CellTrics and incubated on ice for 30 minutes.

The entire suspension was carefully layered onto 0.5 mL of 1.2 M sucrose solution (see Single-nucleus suspension, LiveNuclei protocol) and centrifuged at 5,000 × g for 20 minutes at 2°C. After centrifugation, both phases were aspirated, and the pellet was resuspended in 1 mL of 1 M sucrose solution. The suspension was then layered onto 500 µL of fresh 1.2 M sucrose solution and centrifuged again at 5,000 × g for 20 minutes at 2°C.

Both phases were carefully aspirated, and the nuclei pellet was resuspended in 1 mL of nuclei wash buffer. The sample was then centrifuged at 1,100 × g for 5 minutes at 2°C. After discarding the supernatant, the pellet was resuspended in 500 µL of fresh nuclei wash buffer and incubated on ice for 30 minutes.

Fluorescently labelled antibodies were then added to the suspension—NeuN (1:500, Merck), PU.1 (1:500, CST), and ACSA-2 (1:200, Miltenyi Biotec)—and incubated overnight on a rotating platform at 4°C. After incubation, the sample was centrifuged at 1,100 × g for 5 minutes at 2°C. The supernatant was carefully removed, and the nuclei pellet was resuspended in 1 mL of nuclei wash buffer.

The suspension was filtered through a 30 µm CellTrics filter and prepared for FANS. Shortly before sorting, 1 µL of Hoechst 33258 was added to the suspension, and PU.1⁺/Hoechst⁺/ACSA-2⁻/NeuN⁻ nuclei were sorted into of Smart-seq3xpress plates containing modified lysis buffer (see Generation of Smart-seq3xpress libraries) using FANS under the same parameters as described above (see Single-cell suspension, LiveCells protocol) (**SFig. 2C**). The full and comprehensive protocol has been deposited on protocols.io (30).

### Generation of Smart-seq3xpress libraries

Libraries were prepared according to the published protocol (6), following the specified parameters with minor modifications. Cells and nuclei isolated from the LiveCells, LiveNuclei, and FixedNuclei protocols were sorted into 384-well plates containing 0.3 µL of lysis buffer, dispensed into each well alongside 3 µL of silicone oil (100 cSt, Sigma-Aldrich) using the Formulatrix Mantis. The lysis buffer for LiveCells and LiveNuclei contained 0.5 mM dNTPs (ThermoFisher Scientific), 0.125 µM oligo dTVN30 (IDT), 5% PEG8000 (Sigma-Aldrich), 0.0125 µL SIRV4 diluted 1:2000 (Lexogen), and 0.4 U RNaseOUT (ThermoFisher Scientific) diluted in nuclease-free water (NFW).

For the FixedNuclei protocol, the lysis buffer was adapted by incorporating 2.4 U/mL Thermolabile Proteinase K (New England Biolabs) and diluting it in TE buffer with a lower EDTA concentration (0.1 M, Invitrogen) instead of NFW. Sorted cells and nuclei in the 384-well plates were promptly stored at −80°C until further processing.

After storage at −80°C, the plates were thawed and immediately incubated at 72°C for 10 minutes, followed by reverse transcription and 15 PCR cycles for pre-amplification, as outlined in the protocol. Post pre-amplification, the cDNA was diluted with 9 µL of nuclease-free water (NFW) and stored at −20°C overnight.

The protocol for plates prepared from the FixedNuclei protocol was slightly modified. Initially, RNA-binding proteins were removed by Proteinase K during incubation at 37°C for 30 minutes. The enzyme was subsequently deactivated by incubating the plates at 56°C for 30 minutes, followed by RNA denaturation at 80°C for 15 minutes. Before adding the reverse transcription mix, the plates were cooled to 4°C, after which the standard protocol was resumed.

Subsequently, 1 µL of diluted cDNA from each well was transferred to prepared plates containing dried index primers (2.82 pmol), followed by tagmentation using 0.002 µL of tagmentase per reaction (TDE1 Tn5, Illumina). Tagmentation was stopped by the addition of SDS (ThermoFisher Scientific) to a final concentration of 0.2%.

Libraries were indexed by adding 5 µL of PCR mix containing Tween-20 (ThermoFisher Scientific) to a final concentration of 0.01% and amplified using 14 PCR cycles. The libraries were then pooled and cleaned using SeraMag (GE) beads (0.7×) in 22% PEG, followed by a second clean-up using SPRIselect (Beckman Coulter) at a 0.65× ratio.

The size of the prepared libraries was assessed using the Fragment Analyzer HS NGS Library kit (Agilent), and concentrations were measured using Qubit. Libraries were sequenced on an Illumina NextSeq 2000, loaded at a calculated concentration of 650 pM, and sequenced using 2 × 109 bp paired-end reads and 2 × 10 bp index reads. For the repeated experiment, 2x159bp was used.

### Data analysis

#### Processing of raw data

FASTQ files, including index reads, were generated from cbcl-files using Illumina’s Bcl-convert (version 4.3.6) without index demultiplexing; all reads were exported as *Undetermined*. Low-quality bases were trimmed from Read 2 using TrimmomaticSE (version 0.39) (31) with parameters “ILLUMINACLIP:SmartSeq3xpressAdapter.fa:2:30:10 LEADING:3 TRAILING:3 SLIDINGWINDOW:4:15 MINLEN:36”. Read identifiers from the filtered Read 2 dataset were extracted and used to retain only corresponding pairs from Read 1 and index reads (Index 1 and Index 2) using seqtk subseq (version 1.3-r106) (32). Filtered files were compressed with pigz (version 2.6) and re-evaluated with FastQC (version 0.11.9) (33).

Obtained files were processed using zUMIs according to the published protocol by Hagemann-Jensen (7), following their specified parameters. Briefly, using zUMIs (version 2.7.9e) (34) using parameters specified in **SFile 1** and conda environment included in zUMIs package with STAR (version 2.7.3a), mouse genome version GRCm38 and annotation gencode.vM8. Several downsampled matrices (1 × 10⁴, 2 × 10⁴, 5 × 10⁴, 7.5 × 10⁴, 1 × 10⁵ read pairs) were generated.

For downstream analysis, we selected a downsampling threshold of 20,000 reads per cell, as the differences between isolation protocols were notable at higher read counts. For the replicated experiment, we applied a downsampling threshold of 2,000 reads per cells for the lower sequencing depth.

### Data filtering

The data were further analysed using the Seurat R package (version 5.0.1) (35). Initially, data from all samples underwent exclusion of genes associated ERCC RNA spike (ERCC-) and spatially inferred RNA velocity (SIRV). Individual isolation protocols were filtered according to percentage of spike (percent.spike), the number of detected genes (nFeature_RNA), the total RNA counts (nCount_RNA), and the proportion of mitochondrial RNA (percent.mt). The cut-offs specific for each type of isolation protocol were the following: LiveCells—percent.spike < 80, nFeature_RNA > 1,000, nCount_RNA > 5,000, UMIfraction > 0.5; LiveNuclei—percent.spike < 75, nFeature_RNA > 600, nCount_RNA > 2, 000, UMIfraction > 0.5; FixedNuclei—percent.spike < 60, nFeature_RNA > 1000, nCount_RNA > 5,000, UMIfraction > 0.5. The cut-offs were determined based on the negative control, which did not contain any sorted cells or nuclei. For repeated experiment, the cut-offs specific for each type of isolation protocol were the following: LiveCells—percent.spike < 17, nFeature_RNA > 450, nCount_RNA > 700; FixedNuclei—percent.spike < 15, nFeature_RNA > 450, nCount_RNA > 500.

### Normalization, Integration, and Clustering

For the initial analysis of all obtained cell populations, data were normalized using SCTransform and integrated with CCAIntegration. Visualization was performed using Uniform Manifold Approximation and Projection (UMAP) based on 50 principal components (PCs). Clustering was conducted using the FindNeighbors and FindClusters functions with a UMAP resolution set to 0.3. Cluster identification was carried out by evaluating the expression of well-established marker genes corresponding to expected cell populations.

For a more focused analysis of microglia, data were filtered based on UMAP clustering to select clusters expressing microglia-specific markers. The filtered dataset was normalized using SCTransform and integrated with CCAIntegration. UMAP visualization was generated using 50 principal components (PCs), followed by clustering with the FindNeighbors and FindClusters functions at a UMAP resolution of 0.85.

During this process, cluster 4 was excluded due to nonstandard cell characteristics (**SFig. 3A**). In the LiveCells protocol, cells in this cluster appeared as doublets, suggesting potential cell aggregates. A scatter plot of forward scatter area (FSC.A) versus forward scatter height (FSC.H) was used to visualize and identify these doublets (**SFig. 2B**). In the LiveNuclei and FixedNuclei protocols, the excluded cluster exhibited abnormally high ribosomal content, which is atypical for nuclei preparations, suggesting cytoplasmic contamination or incomplete nuclear isolation (**SFig. 2C–D**).

After excluding cluster 4, the microglia data were reprocessed using SCTransform and reintegrated with CCAIntegration. UMAP visualization was again generated using 50 PCs, followed by clustering with FindNeighbors and FindClusters at a UMAP resolution of 0.8. The refined microglia dataset was then used for all downstream analyses.

### Identification of cluster markers

Normalized and scaled RNA assay data were used to identify marker genes in both the full dataset and the microglia (MG) subset using the FindAllMarkers function with the default Wilcoxon test. Significant markers were defined as those expressed in at least 80% of cells within a cluster (**STab. 1-2**).

For all cell populations, subcluster identities were assigned by comparing the identified markers to known gene signatures from previously described cellular subtypes. Similarly, in the focused MG analysis, subclusters were annotated using established microglial gene signatures from Sala Frigerio et al. (13). Manual curation was applied in both cases to ensure accurate identification and characterization of the cellular subtypes.

The AddModuleScore function in Seurat was used to calculate module scores based on the top 10 marker genes for microglial subtypes from Sala Frigerio et al. (13). The results were visualized using FeaturePlot to display the spatial distribution of subtype-specific gene expression across the UMAP embedding.

### Differential Expression Gene (DEG) Analysis

DEG was performed using Seurat’s FindMarkers function with the default Wilcoxon rank-sum test on normalized and scaled RNA assay data. Comparisons were made between specific clusters, particularly activated response microglia (ARM) and homeostatic microglia (HM), separately for each condition to identify significantly upregulated or downregulated genes. Genes were considered differentially expressed if they were detected in at least 10% of cells in either group, with a log2 fold change (log2FC) > 1 and an adjusted p-value < 0.05 to account for multiple testing (**STab. 3)**.

### Scatterplots

Mean normalized gene expression values were calculated for ARM cells across conditions. A scatter plot was generated to compare gene abundance between the LiveCells protocol (x-axis) and either the FixedNuclei or LiveNuclei isolation protocols (y-axis). Full results are provided (**STab. 4–5)**.

### Gene Ontology Analysis (GO)

GO enrichment analysis was conducted using the compareCluster function from the clusterProfiler R package (version 4.4.4) (36). The analysis was performed on genes with a log-normalized mean expression greater than 1 in both LiveCells and FixedNuclei datasets. The Biological Process (BP) ontology was selected for functional annotation. A significance threshold of adjusted p-value (padj) < 0.05 was applied to identify significantly enriched GO terms. This approach allowed for the comparison of functional pathways between the two conditions, highlighting key biological processes associated with each dataset **(STab. 6)**.

### Cell–cell communication analysis

Cell–cell communication was analyzed using the CellChat R (v1.6.1) package on normalized and scaled RNA assay data (37). Preprocessed CellChat objects for LiveCells, LiveNuclei, and FixedNuclei were loaded and merged using mergeCellChat. Functional network similarities were computed with compute NetSimilarityPairwise and visualized using netEmbedding. Clustering was performed with netClustering to identify similar signaling patterns. The final merged object was saved for further analysis.

## LIST OF ABBREVIATIONS

scRNA-seq: single-cell RNA sequencing
snRNA-seq: single-nucleus RNA sequencing
LiveCells: single whole-cell RNA sequencing without formaldehyde fixation of tissue
LiveNuclei: single-nucleus RNA sequencing without formaldehyde fixation
FixedNuclei: new established protocol - single-nucleus RNA sequencing with formaldehyde fixation
MCAO: middle cerebral artery occlusion
MG: microglia
MON: monocytes
PVM: perivascular macrophages
OL: oligodendrocytes
AST: astrocytes
MUR: mural cells
NEUT: neutrophils
EC: epithelial cells
HM1: homeostatic 1
HM2: homeostatic 2
INTER: intermediate
ARM: activated response microglia
IRM: interferon response microglia
DIV: dividing
GO: gene ontology analysis
UMAP: uniform manifold approximation and projection

## SUPPLEMENTARY FILES

Supplementary files have been included in a separate file.

