## Supplemental figures, file and tables for "PU.1-driven enrichment enables microglia profiling from frozen brain tissue using the high-throughput Smart-seq3xpress method": SFig 1_3.pdf

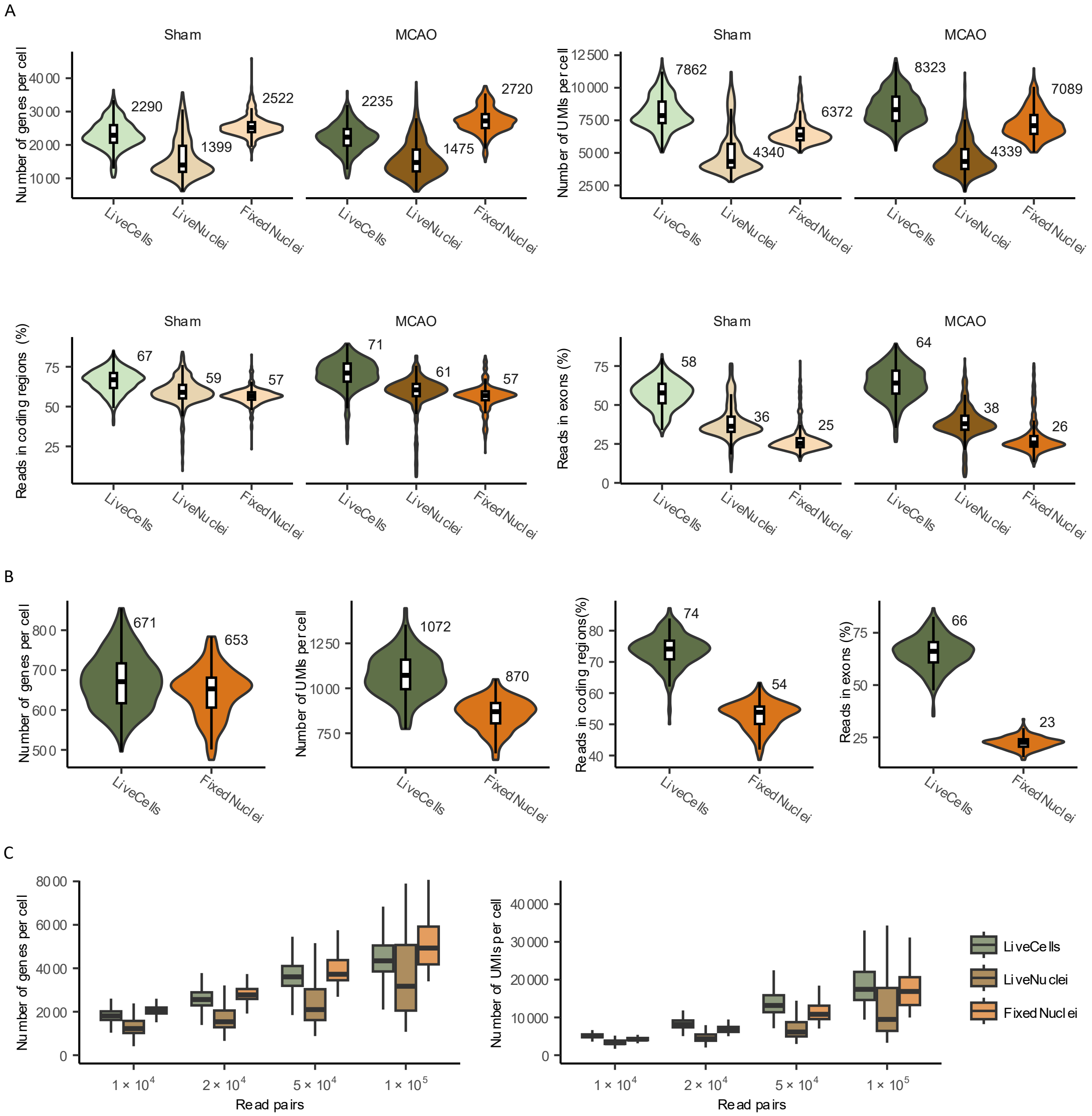


**SFig. 1: Technical parameters for Sham and MCAO condition separately**

**A** Violin plots with overlaid box plots summarizing technical parameters: number of genes per cell, number of UMIs per cell, reads in coding regions and reads in exons for LiveCells, LiveNuclei, and FixedNuclei and compared for Sham and MCAO. Median values are displayed in the top-right corner of each plot. Box plots show medians (black line), first and third quartiles (white boxes), and whiskers extending to 1.5× the interquartile range. **B** Violin plots with overlaid box plots summarizing technical parameters: number of genes per cell, number of UMIs per cell, reads in coding regions and reads in exons for LiveCells, and FixedNuclei for MCAO. Median values are displayed in the top-right corner of each plot. Box plots show medians (black line), first and third quartiles (white boxes), and whiskers extending to 1.5× the interquartile range. **C** Comparison of detected genes and UMIs per cell across varying average reads per cell and isolation protocols (LiveCells, LiveNuclei, and FixedNuclei), shown as box plots (median: black line; quartiles: boxes; whiskers: 1.5× the interquartile range).

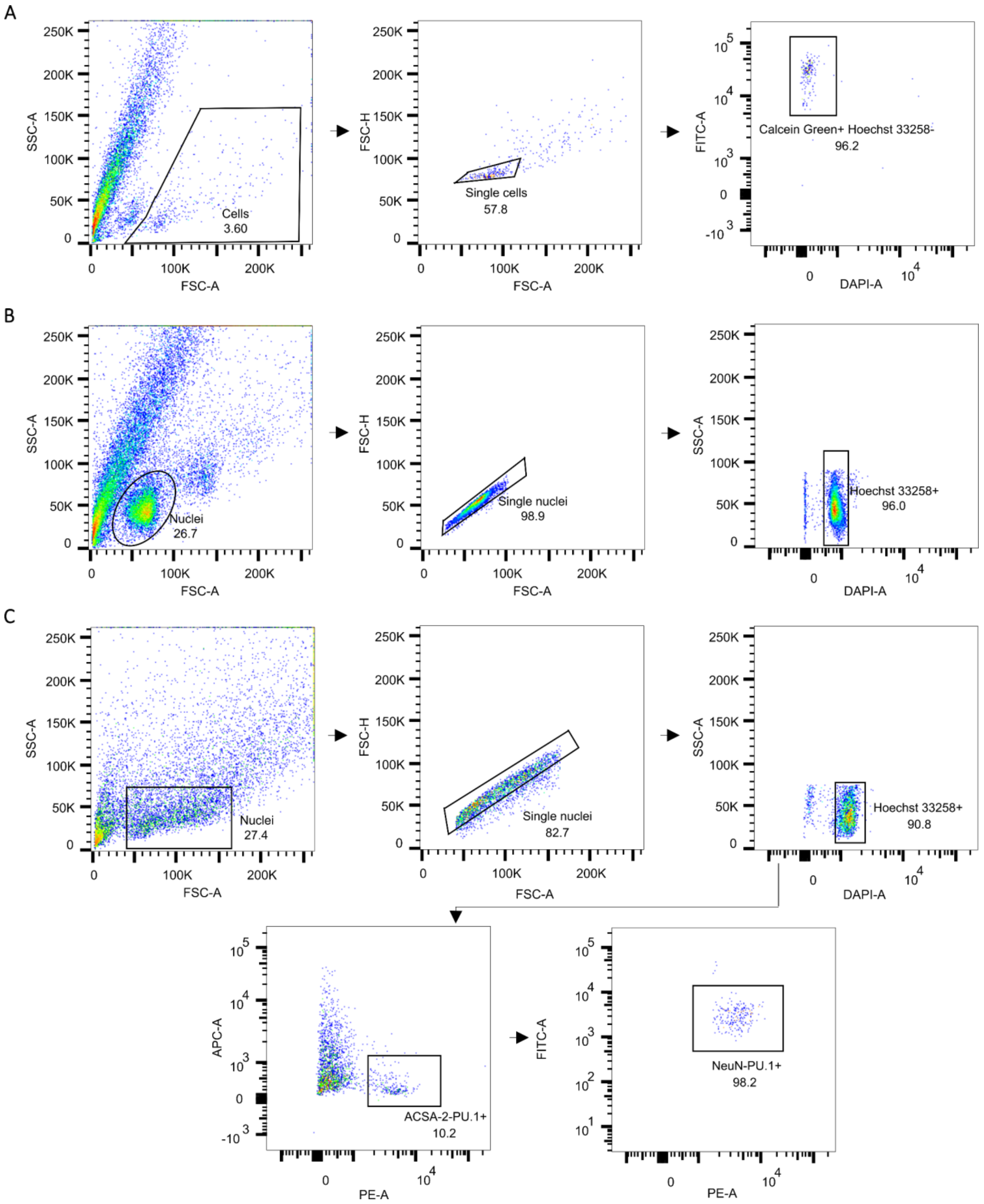


**SFig. 2: Gating strategy for sorting across protocols**

The initial gating for all protocols involved forward scatter area (FSC-A) versus side scatter area (SSC-A) to identify cells or nuclei based on size and granularity. Doublets and aggregates were excluded using FSC-A versus forward scatter height (FSC-H) gating to ensure the selection of singlecells or -nuclei. **A** LiveCells – CD11b+ cells, enriched via MACS, were gated based on characteristic scatter patterns. Live cells were identified as CalceinGreen$^{+}$/Hoechst 332258$^{-}$. **B** LiveNuclei – CD11b+ cells enriched by MACS underwent nuclei isolation. The nuclei were gated based on scatter properties, and Hoechst 332258$^{+}$ nuclei were selected. **C** FixedNuclei – Nuclei were gated using scatter characteristics, and Hoechst 332258$^{-}$/ACSA-2$^{-}$/NeuN$^{-}$/PU.1$^{+}$ nuclei were selected. Data acquisition was performed using FACS, and analysis was completed using FlowJo software (v10.9.0).

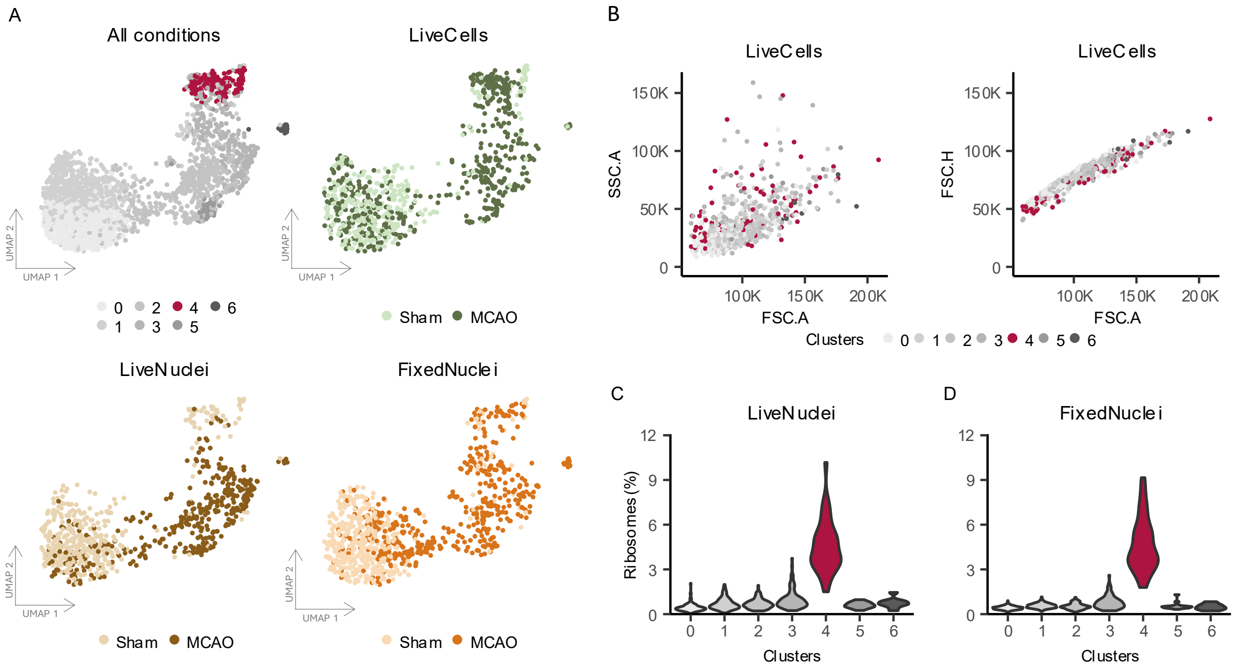


**SFig. 3: Exclusion of cluster**

**A** UMAP visualization showing identified microglia clusters across all conditions with highlight of cluster which was removed for downstream analysis. Additionally, the cluster representation is displayed for each isolation protocol (LiveCells, LiveNuclei, and FixedNuclei) in both Sham and middle cerebral artery occlusion (MCAO) conditions. **B** Scatter plot visualization of fluorescence-activated cell sorting data for LiveCells. The left panel displays FSC.A (forward scatter area) versus SSC.A (side scatter area), while the right panel shows FSC.A versus FSC.H (forward scatter height). Points are color-coded by cluster assignment, highlighting distinct cellular populations. **C** Violin plots illustrating the percentage of ribosomal gene expression across clusters for the LiveNuclei isolation protocol. **D** Violin plots illustrating the percentage of ribosomal gene expressions across clusters for the FixedNuclei isolation protocol.
