## Supplemental figures, file and tables for "PU.1-driven enrichment enables microglia profiling from frozen brain tissue using the high-throughput Smart-seq3xpress method": SFile 1 Smartseq3xpress_SS3x_manuscript_all.yaml

```
project: Smartseq3xpress
sequence_files:
  file1:
    name: /storage/abaffyp/20240802_SS3x_NovaSeqX_mix/raw/Undetermined_S0_L003_R1_001.fastq.gz
    base_definition:
    - cDNA(25-108)
    - UMI(12-21)
    find_pattern: ATTGCGCAATG;2
  file2:
    name: /storage/abaffyp/20240802_SS3x_NovaSeqX_mix/raw/Undetermined_S0_L003_R2_001.fastq.gz
    base_definition: cDNA(1-108)
  file3:
    name: /storage/abaffyp/20240802_SS3x_NovaSeqX_mix/raw/Undetermined_S0_L003_I1_001.fastq.gz
    base_definition: BC(1-10)
  file4:
    name: /storage/abaffyp/20240802_SS3x_NovaSeqX_mix/raw/Undetermined_S0_L003_I2_001.fastq.gz
    base_definition: BC(1-10)
reference:
  STAR_index: /storage/abaffyp/Genomes/GRCm38_SS3x_noGTF
  GTF_file: /storage/abaffyp/Genomes/GRCm38_SS3x_noGTF/gencode.vM8.annotation.gtf
  additional_STAR_params: --clip3pAdapterSeq CTGTCTCTTATACACATCT
  additional_files: /storage/abaffyp/Genomes/SIRV/SIRV_ERCC_longSIRV_multi-fasta_20210507.fasta
out_dir: /storage/abaffyp/20240802_SS3x_NovaSeqX_mix/zUMIs/
num_threads: 32
mem_limit: 512
filter_cutoffs:
  BC_filter:
    num_bases: 4
    phred: 20
  UMI_filter:
    num_bases: 3
    phred: 20
barcodes:
  barcode_num: null
  barcode_file: /storage/abaffyp/20240802_SS3x_NovaSeqX_mix/barcodes_SS3x.txt
  automatic: no
  BarcodeBinning: 3
  demultiplex: no
  nReadsperCell: 100
  discardReads: yes
counting_opts:
  introns: yes
  downsampling: 0,10000,20000,30000,40000,50000,75000,100000,150000,200000,250000,300000,400000,500000,1000000
  strand: 1
  Ham_Dist: 2
  velocyto: yes
  primaryHit: yes
  twoPass: no
make_stats: yes
which_Stage: Filtering
samtools_exec: samtools
pigz_exec: pigz
STAR_exec: STAR
Rscript_exec: Rscript
zUMIs_directory: /home/abaffyp/zUMIs
```
